# Identification of SARS-CoV2-mediated suppression of NRF2 signaling reveals a potent antiviral and anti-inflammatory activity of 4-octyl-itaconate and dimethyl fumarate

**DOI:** 10.1101/2020.07.16.206458

**Authors:** David Olagnier, Ensieh Farahani, Jacob Thyrsted, Julia B. Cadanet, Angela Herengt, Manja Idorn, Alon Hait, Bruno Hernaez, Alice Knudsen, Marie Beck Iversen, Mirjam Schilling, Sofie E. Jørgensen, Michelle Thomsen, Line Reinert, Michael Lappe, Huy-Dung Hoang, Victoria H. Gilchrist, Anne Louise Hansen, Rasmus Ottosen, Camilla Gunderstofte, Charlotte Møller, Demi van der Horst, Suraj Peri, Siddarth Balachandran, Jinrong Huang, Martin Jakobsen, Esben B. Svenningsen, Thomas B. Poulsen, Lydia Bartsch, Anne L. Thielke, Yonglun Luo, Tommy Alain, Jan Rehwinkel, Antonio Alcamí, John Hiscott, Trine Mogensen, Søren R. Paludan, Christian K. Holm

**Author notes:** Correspondence to: Christian Kanstrup Holm, and David Olagnier. These authors contributed equally (Ensieh Farahani, Jacob Thyrsted, Julia B. Cadanet, Angela Herengt). These authors contributed equally (Trine Mogensen, Søren R. Paludan).

## Abstract

Antiviral strategies to inhibit Severe Acute Respiratory Syndrome Coronavirus 2 (SARS-CoV2) and the pathogenic consequences of COVID-19 are urgently required. Here we demonstrate that the NRF2 anti-oxidant gene expression pathway is suppressed in biopsies obtained from COVID-19 patients. Further, we uncover that NRF2 agonists 4-octyl-itaconate (4-OI) and the clinically approved dimethyl fumarate (DMF) induce a cellular anti-viral program, which potently inhibits replication of SARS-CoV2 across cell lines. The anti-viral program extended to inhibit the replication of several other pathogenic viruses including Herpes Simplex Virus-1 and-2, Vaccinia virus, and Zika virus through a type I interferon (IFN)-independent mechanism. In addition, induction of NRF2 by 4-OI and DMF limited host inflammatory responses to SARS-CoV2 infection associated with airway COVID-19 pathology. In conclusion, NRF2 agonists 4-OI and DMF induce a distinct IFN-independent antiviral program that is broadly effective in limiting virus replication and suppressing the pro-inflammatory responses of human pathogenic viruses, including SARS-CoV2.

**One Sentence Summary:** NRF2 agonists 4-octyl-itaconate (4-OI) and dimethyl fumarate inhibited SARS-CoV2 replication and virus-induced inflammatory responses, as well as replication of other human pathogenic viruses.

## Introduction

The 2020 SARS-CoV2 pandemic emphasizes the urgent need to identify cellular factors and pathways that can be targeted by new broad-spectrum anti-viral therapies. Viral infections usually cause disease in humans through both direct cytopathogenic effects and through excessive inflammatory responses of the infected host. This also seems to be the case with SARS-CoV2 as COVID-19 patients develop cytokine storms that are very likely to contribute to, if not drive, immunopathology and severe disease(*1, 2*). For these reasons, anti-viral therapies must aim to not only inhibit viral replication but also to limit inflammatory responses of the host. Nuclear factor (erythroid-derived 2) - like 2 (NRF2) functions as a cap’n’collar basic leucine zipper family of transcription factors characterized structurally by the presence of NRF2-ECH homology domains(*3*). At homeostasis, NRF2 is kept inactive in the cytosol by its inhibitor protein KEAP1 (Kelch-like ECH-associated protein 1), which targets NRF2 for proteasomal degradation(*4*). In response to oxidative stress, KEAP1 is inactivated and NRF2 is released to induce NRF2-responsive genes. In general, the genes under the control of NRF2 protect against stress-induced cell death and NRF2 has thus been suggested as the master regulator of tissue damage during infection(*5*). Importantly, NRF2 is now demonstrated as an important regulator of the inflammatory response(*6, 7*) and functions as a transcriptional repressor of inflammatory genes, most notably interleukin (IL-) 1β, in murine macrophages(*8*). Recent reports have now demonstrated that NRF2 is induced by several cell derived metabolites including itaconate and fumarate, to limit inflammatory responses to stimulation of TLR signaling with lipopolysaccharide stimulation(*9*). The chemically synthesized and cell-permeable derivative of itaconate, 4-octyl-itaconate (4-OI) was then demonstrated to be a very potent NRF2 inducer(*9*). Of special interest is the derivative of fumarate, dimethyl fumarate (DMF), a US Food and Drug Administration (FDA) approved drug, which is used as an anti-inflammatory therapeutic in multiple sclerosis (MS) and demonstrated, at least in animal models, a potent capacity to suppress pathogenic inflammation through a Nrf2-dependent mechanism(*10, 11*). Besides limiting the inflammatory response to LPS, induction of NRF2 by 4-OI also inhibits the Stimulator of Interferon Genes (STING) antiviral pathway along with interferon (IFN) stimulated gene expression(*12*). In opposition to this anti-viral effect of NRF2 on the IFN-response a recent single-cell RNA-seq analysis has demonstrated that NRF2 gene expression signatures correlated negatively with susceptibility to HSV1 infection (*13*). If NRF2 agonists can be used to limit viral replication of SARS-CoV2 or other pathogenic viruses is, however, not known.

Here we demonstrate that expression of NRF2-dependent genes is suppressed in biopsies from COVID-19 patients and that treatment of cells with NRF2 agonists 4-OI and DMF induces a strong anti-viral program that limits SARS-CoV2 replication. The anti-viral effect of activating NRF2 extended to other pathogenic viruses including Herpes Simplex Virus-1 and-2 (HSV-1 and HSV-2), Vaccinia Virus (VACV), and Zika Virus (ZIKV). Further, 4-OI and DMF limited the release of pro-inflammatory cytokines in response to SARS-CoV2 infection and to virus-derived ligands through a mechanism that limits IRF3 dimerization. In summary, we demonstrate that NRF2 agonists are plausible broad-spectrum anti-viral and anti-inflammatory agents and we suggest a repurposing of the already clinically approved DMF for the treatment of SARS-CoV2.

## Results

### NRF2 dependent anti-oxidant response is suppressed in COVID-19 patient biopsies

To identify host factors or pathways that are important for controlling SARS-CoV2 infection, publicly available transcriptome data sets including transcriptome analysis of lung biopsies from COVID-19 patients were analyzed using differential expression analysis(*14*). Here, genes linked with inflammatory and anti-viral pathways, including RIG-I receptor and Toll-like receptor signaling, were highly enriched in COVID-19 lung patient samples, whereas genes associated with the NRF2 dependent anti-oxidant response were highly suppressed (**Fig. 1a-c)**. That NRF2-induced genes are repressed during SARS-CoV2 infections was supported by re-analysis of another data-set builing on transcriptome analysis of lung autopsies obtained from five individual COVID-19 patients (Desai et al., 2020) (**Fig. 1d**). Further, that the NRF2-pathway is repressed during infection with SARS-CoV2 was also supported by *in vitro* experiments where the expression of NRF2-inducible proteins Heme Oxygenase 1 (HO-1) and NAD(P)H quinone oxydoreducatse 1 (NqO1) was repressed in SARS-CoV2 infected Vero hTMPRSS2 cells while the protein levels of canonical anti-viral transcription factors such as STAT1 and IRF3 seemed unaffected **(Fig. S1)**. These data indicate that SARS-CoV2 targets the anti-oxidant NRF2 pathway and thus suggests that the NRF2 pathway restricts SARS-CoV2 replication.

**Fig. 1.**
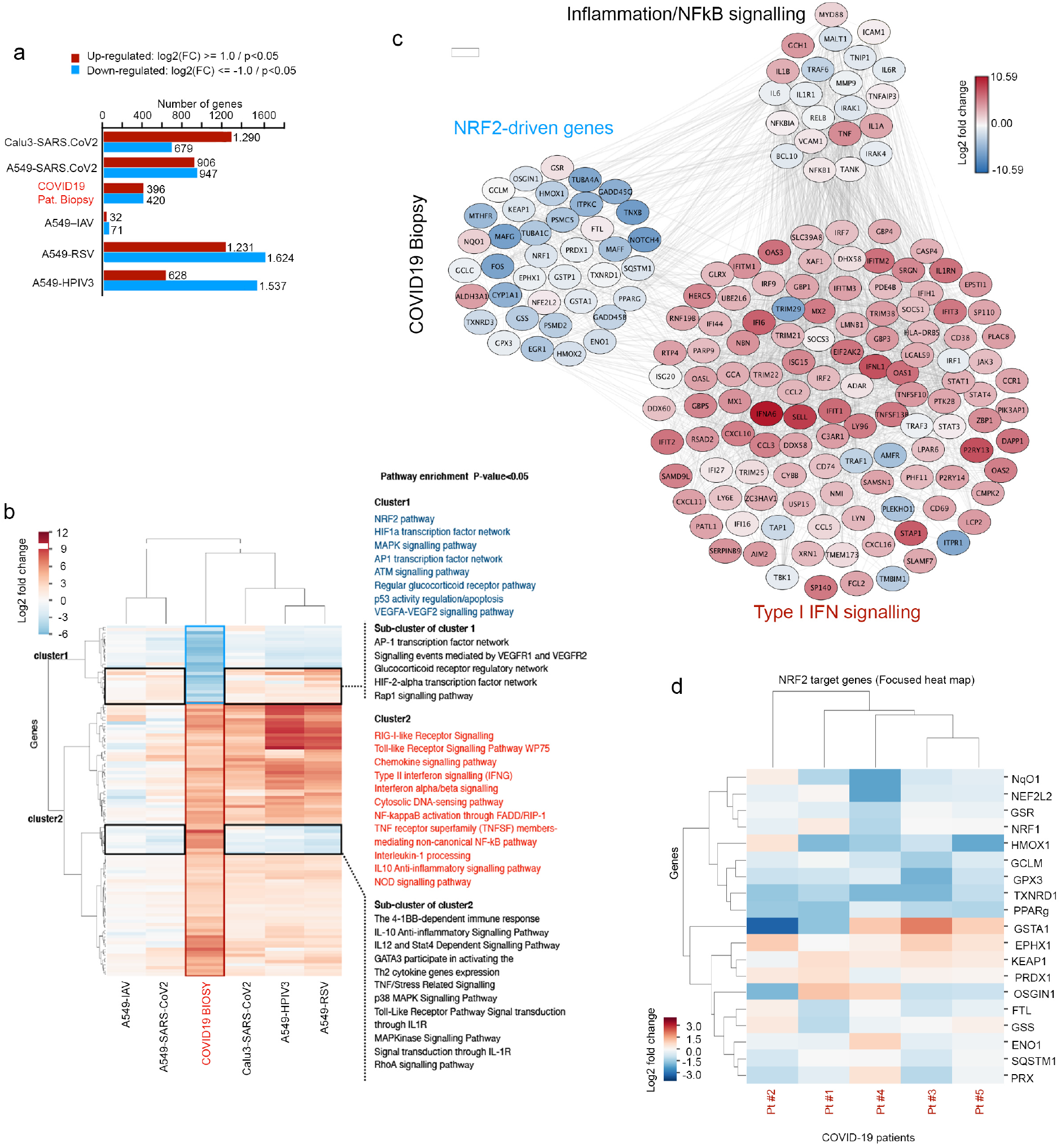
Expression of NRF2-driven genes is suppressed in COVID-19 patient biopsies. **(a, b, and c)** Reanalysis of data published by Blanco-Melo *et al*., 2020. **(a)** Bar-chart of the number of transcripts that show differential expression (up and down). Genes with p<0.05 and Log2 fold change in COVID-19 patient lung biopsies were normalized against healthy lung biopsies, and in cell lines Calu3, NHBE and A549 infected with either SARS-CoV2, Influenza A virus (IAV), Respiratory Syncytial virus (RSV), or human parainfluenza virus type 3 (HPIV3) were normalized against mock treated cells. **(b)** Cloud analysis of NRF2-driven differentially expressed genes. Subsets annotated as inflammation/NFκB signaling and Type I IFN signaling exhibit different expression patterns. The experiment is a re-analysis of data from Blanco-Melo *et al*., https://doi.org/10.1101/2020.03.24.004655. **(c)** Heat map of the subset of genes significantly differentially expressed in COVID19 biopsies and simultaneously differentially expressed in at least 3 of the other conditions tested. The genes in each cluster were used for pathway enrichment analysis. Genes in cluster 1 are dominantly down-regulated in COVID19 biopsies while a sub-cluster of genes in cluster 1 are up-regulated in the cell lines. Conversely, genes in cluster 2 are predominantly up-regulated in biopsies and in most other test-samples. A sub-cluster of the genes in cluster 2 are down regulated in the cell lines. For each cluster, the significantly enriched pathways are listed (EnrichR). **(d)** Reanalysis of the data from Desai N et al. (May 2020), GEO accession code GSE150316. Heat map of NRF2 target genes of the lung autopsies from the five COVID19 patients. Healthy lung samples were used as negative control.

### NRF2 agonists 4-OI and dimethyl fumarate are strong inhibitors of SARS-CoV2 replication

Considering that NRF2 suppresses anti-viral IFN-responses, it was surprising to discover that treatment of Vero cells with 4-OI generated before infection with SARS-CoV2 (strain #291.3 FR-4286 isolated from a patient in Germany) resulted in a remarkable reduction in SARS-CoV2 RNA levels in a dose-dependent manner **(Fig. 2a+b)** while not affecting cell viability as determined by lactate dehydrogenase (LDH) assay **(Fig. S2)**. Further, subsequent release of progeny SARS-CoV2 virus particles to the cell supernatant was equally decreased by 4-OI treatment as measured by TCID50 assay to quantify virus by dilution of virus-induced cytopathogenic effects and by plaque assay **(Fig. 2c-f)**. The reduced viral replication led to reduced virus-induced cytotoxicity of the infected Vero cells determined by lactate dehydrogenase release assay and by immunoblotting for cleaved Caspase 3 and Poly(ADP-Ribose) Polymerase 1 (PARP-1), which are hallmark indicators of apoptosis(*15*) **(Fig. 2g+h)**. Interestingly, the observation that the NRF2 pathway is inhibited in response to SARS-CoV-2 infection could be recapitulated in SARS-CoV-2 infected Vero cells as demonstrated by both the basal decrease in NRF2-driven proteins HO-1 and NqO1 and their incapacity to be induced by Nrf2 agonist 4-OI (**Fig. 2h**). The effect of 4-OI was also retained in the lung cancer cell line Calu-3, where SARS-CoV2 RNA levels were reduced by >2-logs **(Fig. 2i),** while release of progeny virus was reduced by > 6-logs based on TCID50 analysis of cell supernatants **(Fig. 2j+k)**. In the immortalized human epithelial cell line NuLi, total infection levels were relatively low compared to what we could observe in Calu3 and Vero cells but 4-OI treatment still reduced SARS-CoV2 RNA levels and release of progeny virus (**Fig. 2l+m)**. We further tested the anti-viral effect towards SARS-CoV2 in primary human airway epithelial (HAE) cultures **(Fig. 2n)**. Here, 4-OI treatment also significantly reduced viral RNA levels **(Fig. 2o)**. Interestingly, when treating Calu3 cells with DMF, another known NRF2 inducer and a clinically approved drug in the first-line-of treatment of multiple sclerosis, we could also observe an anti-viral effect toward SARS-CoV2 replication similar in magnitude as what we had observed with 4-OI **(Fig 2p-q**) as well as a reduced but significant effect when using Vero cells **(Fig. 2r)**. To further evaluate if the NRF2/KEAP1 axis controls an anti-viral pathway effective in inhibiting SARS-CoV2 replication, we used genetic activation of NRF2 by silencing of KEAP1. This approach supported the 4-OI data as silencing of KEAP1 led to suppressed replication of SARS-CoV2 in Calu3 cells by qPCR analysis, immunoblotting, and TCID50 analysis **(Fig. 2s-u)**. Finally, the anti-viral effect of SARS-CoV2 was not isolated to this particular isolate as the effect of 4-OI was reproduced using a different SARS-CoV2 isolate obtained in Japan(*16*) **(Fig. 2v+x)**. These data demonstrate that NRF2 inducers 4-OI and DMF induce potent anti-viral responses that efficiently inhibit SARS-CoV2 replication across multiple cellular systems.

**Fig. 2.**
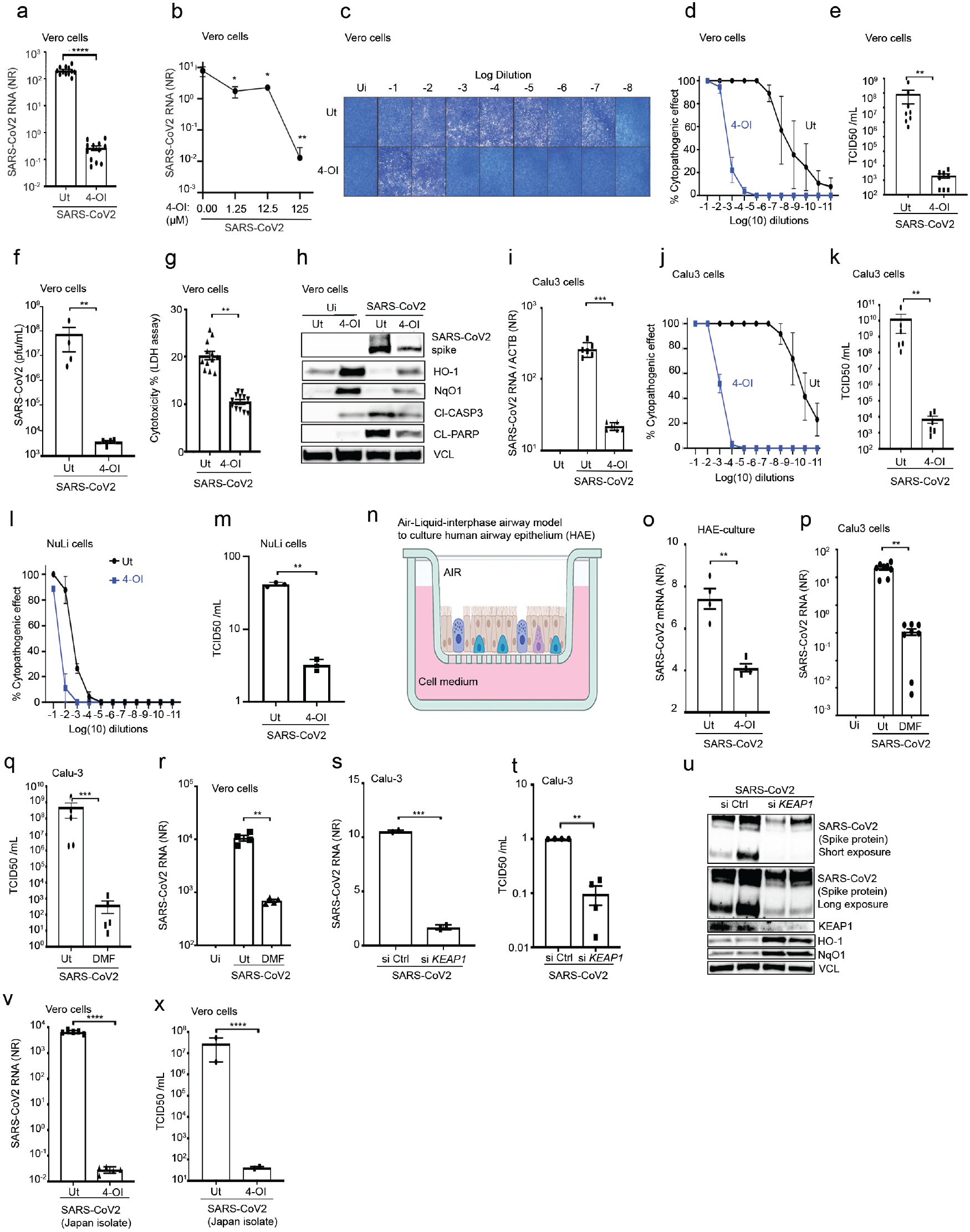
4-Octyl-itaconate (4-OI) and dimethyl fumarate (DMF) inhibit SARS-CoV2 replication. **(a-e)** Vero E6 expressing human TMPRSS2 cells were treated with 4-OI (125μM) or lower concentrations were indicated for 48h and subsequently infected with SARS-CoV-2 at a MOI of 0.1 for 48h, respectively. Infection and replication were assessed by qPCR and TCID50 or plaque assay, respectively. Data displayed in a and e are means and s.e.m. of pooled data from three independent experiments with each data-point representing one biological sample. Data displayed in b, c, and d are represenatative of two independent experiments and indicating means and s.e.m.. In **(f),** supernatants from (e) were also assessed for viral titers by semi-solid agarose plaque assay. Data are pooled from two independent experiments with each data-point representing one biological sample. **(g)** Vero E6 expressing human TMPRSS2 cells were treated with 4-OI (125μM) for 48h and subsequently infected with SARS-CoV-2 at a MOI of 0.1 for 48h. Cell death was evaluated using an LDH release assay. Data are pooled from two independent experiments each performed in sextuplicate displaying the means and s.e.m. In **(h),** whole cell extracts from (g) were immunoblotted for various Nrf2 and death associated markers. Blot is representative of two independent experiments. In **(i-k)** Calu3 cells were treated with 4-OI (100μM) for 48h and subsequently infected with SARS-CoV-2 (MOI 0.5) for 48h. Viral RNA content and progeny virus were assessed by qPCR and TCID50, respectively. Data are the means and s.e.m. of pooled data from two independent experiment performed in triplicate. In **(l-m),** NuLi cells were treated with 4-OI (100μM) for 48h and subsequently infected with SARS-CoV-2 (MOI 0.5) for 48h. Viral titers were determined by TCID50. Data are the means and s.e.m. and representative of two independent experiments performed in duplicate. In, **(n)** schematic of an HAE culture. **(o)** HAE cultures (n=4) were treated with 4-OI (125μM) overnight and were subsequently infected with SARS-CoV-2 (MOI 0.1). Viral genome content was assessed by qPCR (mean +/-s.e.m. from 4 independent primary HAE cultures). In **(p-r),** Calu-3 cells and Vero E6 expressing hTMPRSS2 cells were treated with dimethyl fumarate (DMF) (150-200 μM) for 48h and subsequently infected with SARS-CoV-2 at a MOI of 0.5 and 0.1, respectively. Data in (p and q) are pooled from two independent experiments with means and s.e.m. In **(s-u)**, Calu-3 cells were treated with indicated siRNAs for 48 hours before infection with SARS-CoV2 at a MOI of 0.5. Cell pellets were then etiher collected for RNA extraction and qPCR analysis (s) or for immunoblotting (u). Cell supernatants were analysed by TCID50 assay (t). Data are mean and s.e.m and representative of two independent experiments. In **(v-w)**, Vero cells were treated with 4-OI at 125μM before infection with a different SARS-CoV2 isolate obtained from Japan(*16*). Data in **u** and **v** are obtained from on experiment performed in duplicate. All statistical analysis were performed using a two-tailed Student’s *t*-test where **p< 0.01, ***p<0.001, and ****p<0.0001.

### Activation of NRF2 with 4-OI broadly inhibits viral replication through an IFN-independent pathway

The anti-viral effect of 4-OI was not restricted to SARS-CoV2 but also extended to other human pathogenic viruses. Using the human keratinocyte cell line HaCaT as a model of Herpes Simplex Virus (HSV) type 1 and 2 we could observe that treatment with 4-OI reduced both the release of progeny virus, the cellular content of virus RNA determined by RNA sequence analysis,and viral protein as determined by both immunoblotting and flow cytometry (**Fig. 3a-e and Fig. S3**). By contrast, the expression of NRF2-inducible HO-1, NqO1, and Sequestosome 1 (SQSTM1) was highly increased in response to 4-OI treatment (**Fig. 2c and Fig. S3**). The anti-viral effect of 4-OI was at least partially dependent on NRF2 as silencing hereof by siRNA clearly reduced the suppression of HSV1 infection by 4-OI (**Fig. 2f-g**).

**Fig. 3.**
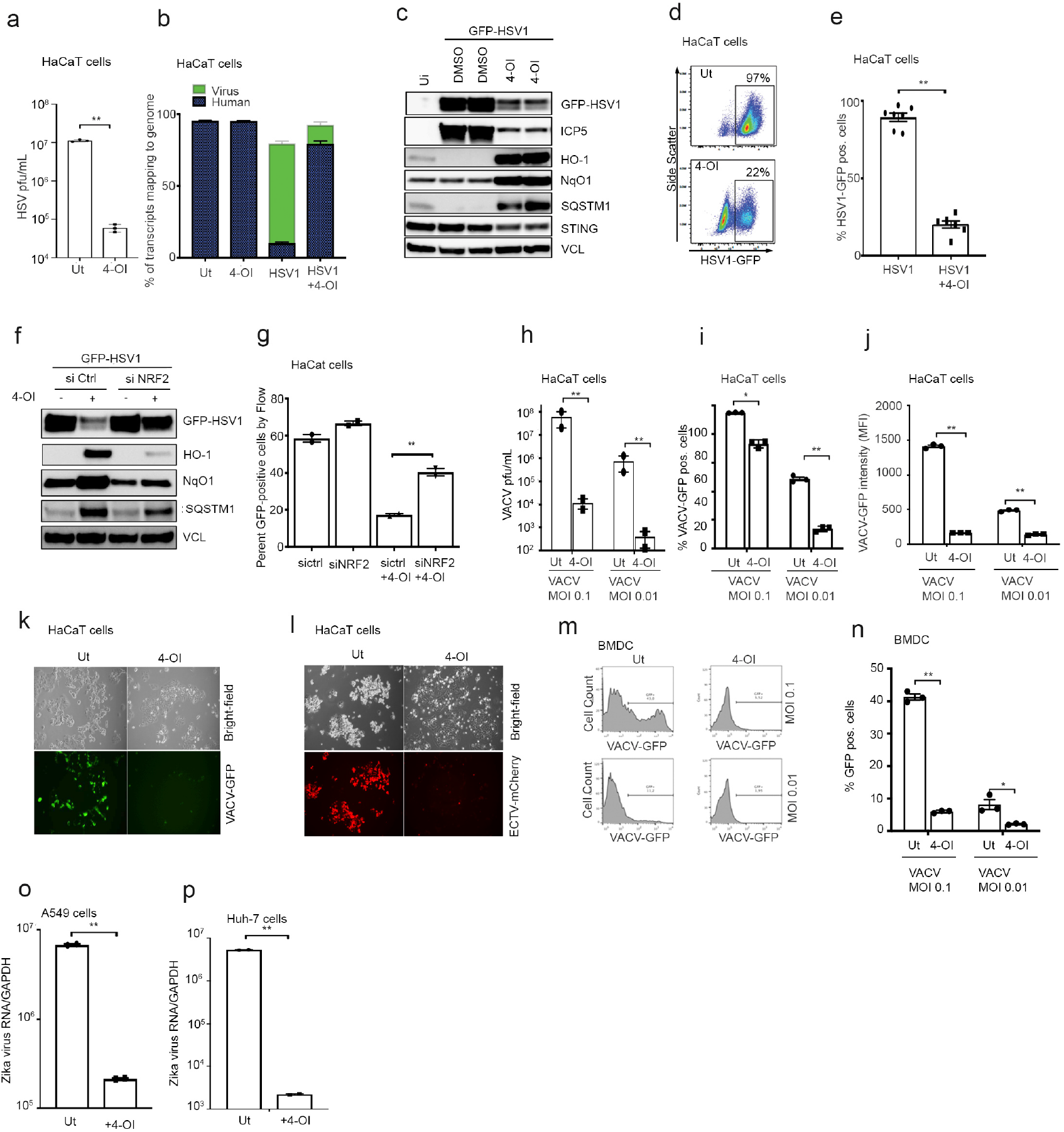
4-OI broadly inhibits other pathogenic viruses including HSV, VACV, and Zika Virus. **(a)** HaCaT cells treated with 4-OI (125μM) for 48h and infected with HSV1-GFP (MOI 0.01). Viral titers were determined by plaque assay. Data are obtained from one experiment of at least seven independent experiments duplicates **(b)** RNA analyzed using RNA-sequencing (n=3) performed once. **(c-e)** HaCaT cells treated with 4-OI (125μM) and infected with HSV1-GFP (MOI 0.01). Lysates were analyzed by immunoblotting with Vinculin (VCL) as loading control and by flow cytometry (n=7 from three independent experiments, mean +/-s.e.m.). **(f-g)** HaCaT cells were lipofected with siRNA for 72h, subsequently challenged with 4-OI (125μM) before HSV1-GFP infection (MOI 0.01) for 24h. Infectivity and silencing efficiency was determined by immunoblotting (f) and flow-cytometry (g). Data obtained from one experiment representative of three independent experiments. **(h-n)** HaCaT cells (h-l) and BMDCs (m-n) were treated with 4-OI (125μM) for 48h and infected with VACV expressing either GFP or ECTV expressing mCherry for 24h. Viral titers and infectivity were determined by plaque assay (h), flow cytometry (i-j and m-n), and visualized by confocal microscopy (k-l). Data are obtained from one experiment representative of two independent experiments. **(o-p)** A549 and Huh-7 cells were pre-treated with 4-OI for 48h (150μM) and infected with Zika virus (ZIKV) (MOI 0.1) for 4 days. Viral genome was determined by qPCR. Data were obtained from one experiment representative of two independent experiments. All statistical analysis were performed using a two-tailed Student’s *t*-test to determine statistical significance where **p< 0.01, ***p<0.001, and ****p<0.0001.

Vaccinia virus (VACV) belongs to the family of human pathogenic poxviruses. We used HaCaT cells, but also bone marrow derived dendritic cells (BMDCs), to test if the anti-viral effect of 4-OI extended to these viruses. Here we could observe that both HaCaT cells and BMDCs became highly resistant to infection with VACV when these were pre-treated with 4-OI as measured by plaque assay and flow cytometry **(Fig. 2h-n)**. This seemed also to be the case for another poxvirus Ectromelia virus (ECTV) as assesses by confocal imaging **(Fig. 2k-l)**. For both HSV1 and VACV the anti-viral effect of 4-OI was extended to other cell type including murine cancer cell line 4T1 and human renal carcinoma 786-O cells **(Fig. S4).** Interestingly, the anti-viral effect of 4-OI was not extended to infection with vesicular stomatitis virus (VSVd51M) emphasizing that the anti-viral program induced by 4-OI effectively inhibits replication of many, but not all, viruses **(Fig. S4)**. The anti-viral effect of 4-OI relied on intracellular restriction of replication, since viral entry was not affected by 4-OI treatment – if anything it seemed to be slightly increased **(Fig. S5)**.

To determine if the anti-viral effect of 4-OI extended to an *in vivo* model of viral pathogenesis, female C57BL6J mice were treated with 4-OI prior to vaginal inoculation with HSV; pre-treatment with 4-OI decreased disease progression **(Fig. S6)**, an effect that was enhanced in mice deficient in STING (*TMEM173*^−/−^) most likely to due to the pro-viral effect 4-OI has on the STING signaling pathway(*12, 17*), which is eliminated in these mice.

Finally, we tested the efficacy of 4-OI on Zika virus, an important human pathogenic virus causing mild symptoms in the competent adult but severe disease when transmitted *in utero*(*18*). Here, we could demonstrate that the anti-viral program induced by 4-OI reduced replication of Zika virus in the human lung cancer cell line A549 and in the human liver cell line Huh-7 **(Fig. 2o-p)**. Given that Vero cells are deficient in type I IFN(*19*), this suggested that the inhibitory effect of 4-OI was actually independent of type I IFN signaling. To address this possibility, we used either HaCaT cells deficient in IFN alpha receptor 2 (IFNAR2), Signal Transducer and Activator of Transcription 1 (STAT1), both of which is necessary for type I IFN-signaling(*20*); or deficient in STING, which is central to type I IFN-response to DNA viruses. Here, cells were treated with 4-OI, followed by infection with HSV1 and VACV. Replication of both viruses was inhibited by 4-OI in STAT1 KO cells and for HSV1 also in STING KO cells as measured by plaque assay and expression of viral proteins by immunoblotting and flow cytometry (**Fig. S7**). In conclusion, 4-OI induces an anti-viral program that operates independently of IFN signaling **(Fig. S7)**.

To examine what general pathways are affected by 4-OI treatment either alone or during infection that could predict the antiviral mode of action itaconate, we treated HaCaT cells with 4-OI before infection with HSV-1. RNA was then collected and analysed by RNA sequencing. Pathway analysis was then used to compare untreated cells to 4-OI treated cells either with or without infection with HSV-1. This analysis identified several pathways that were either induced or repressed by 4-OI treatment while re-confirming that IFN-signaling pathways was repressed by 4-OI **(Fig. S8)**. Amongst the top up-regulated genes induced by 4-OI was the heme oxygensase-1, an enzyme canonicaly involved in stress detoxification, also reported to have antiviral activity against amongst others Zika, Dengue and Ebola viruses (*21–23*). To assess whether, HO-1 had any antiviral activity in our cellular system, Vero hTMPRSS2 and Calu-3 cells were either transfected with an overexpression plasmid encoding HO-1 or genetically silenced for KEAP1 and HO-1 by siRNA, respectively before infection with SARS-CoV-2. None of the treatments (HO-1 overexpression or silencing) really altered SARS-CoV-2 infection/replication suggesting of an HO-1-independent antiviral program induced by NRF2 (**Fig. S9**).

In an attempt to pin-point the anti-viral mode of action of 4-OI, we also used microscope-based analysis of morphology by cell-paint technology **(Fig. S10)**. With this analysis we are able to compare morphological changes in cells treated with 4-OI to cells treated with compounds that have known cellular targets and with cells treated with other compounds with reported anti-viral activity towards SARS-CoV2 including Remdesivir and Hydroxychloroquine(*24, 25*). In this analysis, 4-OI was determined to have an low but significant morphological activity whitout loss of cell viability. Interestingly, the activity of 4-OI did not seem to overlap with other compound with known perturbation in cell morphology including Rapamycin, Bafilomycin, Tunicamycin, Cyclohexamide, Emetine, Mitomycin, or Doxorubicin. Interestingly, there was also no observable overlab with the activity profile of Remdesivir or hydroxychloroquine indicating that the anti-viral mode of action of 4-OI is distinct from known anti-viral mechanisms.

### 4-OI and DMF suppress the inflammatory response to SARS-CoV2

In COVID-19, an uncontrolled pro-inflammatory cytokine storm contributes to disease pathogenesis and lung damage (*26*). For this reason, we investigated if 4-OI and DMF could inhibit expression of pro-inflammatory cytokines induced by SARS-CoV2. In Calu-3 cells, infection with SARS-CoV2 increased the expression of *IFNB1*, C-X-C motif chemokine 10 (*CXCL10*), Tumor Necrosis Factor alpha (*TNFA*), *IL-1B* and C-C chemokine ligand 5 (*CCL5*). Interestingly, this was abolished by pretreatment with 4-OI thus severely reducing the pro-inflammatory response to SARS-CoV2 **(Fig. 4a-b)**. By contrast, expression of the NRF2 inducible gene HMOX1 was highly increased in response to 4-OI treatment **(Fig. 4c)**. The potential anti-inflammatory effect of 4-OI in this context was supported when using HAE cultures. Here, treatment with 4-OI also reduced the expression of *IFNB1*, *CXCL10*, *TNFα*, and CCL5 in the context of SARS-CoV2 infection **(Fig. 4d-e)**, while increasing the expression of the NRF2 inducible gene *HMOX1* **(Fig. 4f)**. A similar pattern was seen in experiments where Calu3 cells were treated with DMF before SARS-CoV2 infection. Here, *IFNB1*, *CXCL10* and *CCL5* mRNA levels were highly reduced in DMF treated cells while *TNFA* mRNA levels seemed unaffected (**Fig. 4g+h**). By contrast, treatment with DMF increased the mRNA expression levels of NRF2 inducible gene *HMOX1* (**Fig. 4i**). As inflammatory responses often stem from immune cells we also tested the effect of 4-OI on Peripheral Blood Mononuclear Cells (PBMCs) harvested from healthy donors. Although stimulation of PBMCs with SARS-CoV2 yielded a very weak induction of *CXCL10* compared to sendai virus (SeV) infection, and no detectable induction of other cytokines, 4-OI treatment also reduced *CXCL10* mRNA levels in this context **(Fig. 4j).** Further, when using PBMCs harvested from four individual patients with severe COVID-19 and admitted to hospital Intensive Care Units (ICUs), we could conclude that in three out of four patients, expression levels of CXCL10 were increased when compared to healthy controls; and that in all four patients, these levels were strongly reduced to or below normal when treating the PBMCs with 4-OI **(Fig. 4k)** indicating that 4-OI is able to relieve inflammatory responses induced by SARS-CoV2 in PBMCs *in vivo*. That CXCL10 levels is a relevant readout in SARS-CoV2 was recently supported by a report by Cheemarla et al., 2020 (https://doi.org/10.1101/2020.06.04.20109306) demonstrating that CXCL10 expression is increased in the upper airways of patients infected with SARS-CoV2.

**Fig. 4.**
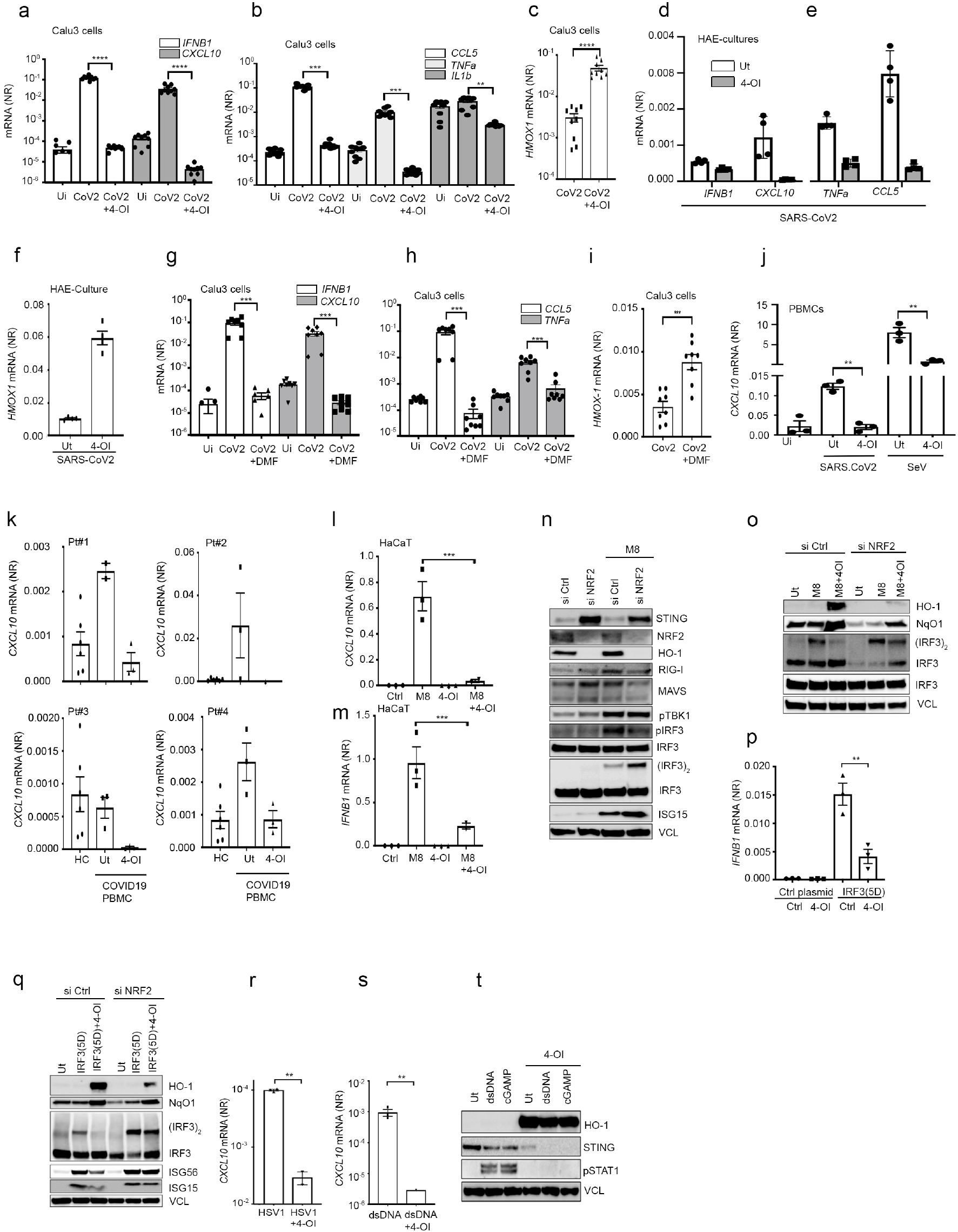
4-OI and DMF limit SARS-CoV2 and HSV induced inflammatory responses. **(a-c)** Calu-3 treated with 4-OI (125μM) for 48h before infection with SARS-CoV2 (MOI 0.5) for 48 hours. RNA was extracted for analysis by qPCR. Graphs display pooled data from three independent experiments with each data-point representing one biological sample where means and s.e.m. are displayed. (**d-f)** HAE cultures (n=4,) were treated overnight with 4-OI at 125μM before SARS-CoV2 infection (MOI 0.1) for 24 and analysis by qPCR. Data are representative of two independent experiments with two donors. **(g-i)** Calu-3 treated with DMF (200 μM) for 48h before infection with SARS-CoV2 (MOI 0.5) for 48 hours. RNA was extracted for analysis by qPCR. In (a+b+c+g+h+i) Graphs display pooled data from three independent experiments with each data-point representing one biological sample where means and s.e.m. are displayed. In (d-f) data are representative of four independent primary HAE cultures where means and s.e.m. are displayed. **(j)** Healthy PBMCs were pre-treated overnight with 4-OI (100μM) before a challenge with SARS-CoV-2 (MOI 10) or sendai virus (SeV) (50 HAU) for an additional 24h. *CXCL10* mRNA levels were determined by qPCR. Data are from one healthy donor in triplicate representative of two independent healthy donors. (**k)** PBMCs from four COVID-19 patients (k) and 2 healthy controls (HC) treated with 4-OI at 100μM overnight before analysis by qPCR. (**l-m)** HaCaT cells were treated with 4-OI (125μM) before stimulation with the sequence optimized RIG-I agonist M8 (10ng/mL) for 6 hours followed by qPCR gene expression analysis. Data represent the means and s.e.m. of one experiment performed in triplicate. **(n)** HaCaT cells were lipofected with indicated siRNAs for 72h before treatment with M8 (10 ng/mL) for 3 hours followed by analysis by immunoblotting. **(o)** HaCaT cells were lipofected with indicated siRNAs for 72h before treatment with 4-OI (125μM) for 48h and stimulation with M8 (10 ng/mL) for 3 hours followed by analysis by immunoblotting. (**p-q)** HEK293(p) and HaCaT(q) cells were transfected with indicated plasmids before treatment with 4-OI at 125μM. In (q), HaCaT cells were lipofected with siRNAs for 72h before plasmid transfection. Cells were then collected for analysis by qPCR (p) and immunoblotting (q). (**r-t)** HaCaT cells were treated with 4-OI at 125μM before infection with HSV1 at MOI 0.01 or transfection with dsDNA (4 μg.mL^-1^). Cell pellets were collected for qPCR and immunoblotting at 6 and 3 hours respectively. In (n-t), data display data from one experiment representative of at least to independent experiments. All statistical analysis were performed using a two-tailed Student’s *t*-test to determine statistical significance where **p< 0.01, ***p<0.001, and ****p<0.0001.

The observed decrease in anti-viral and pro-inflammatory responses could possibly be explained by the 4-OI mediated reduction in cellular viral RNA with subsequent reduced induction of cytokines through cellular RNA sensors such as RIG-I. We therefore investigated the effect of 4-OI on the induction of IFN and of IFN stimulated genes (ISGs) responses activated by a sequence optimized RIG-I agonist M8(*27*). Interestingly, 4-OI treatment reduced IFN-responses induced by this RIG-I agonist M8 **(Fig 4l-m)**, through an effect linked to the inhibition of Interferon Regulatory Factor 3 (IRF3) dimerization but not of upstream phosphorylation of Tank Binding Kinase 1 (TBK1) or of IRF3 expression itself **(Fig 4n)**. Importantly, NRF2 expression itself was closely associated with the inhibition of IRF3 dimerization and host antiviral gene expression, since NRF2 silencing by siRNA was sufficient to restore IRF3 dimerization and limit the inhibitory effect of 4-OI **(Fig. 4n-o)**. When using the constitutively active form of IRF3, IRF3(5D) (*28*), 4-OI was still able to block IRF3 dimerization, and again, this effect was eliminated when NRF2 expression was suppressed by siRNA **(Fig 4p-q)**. These data indicate that an NRF2 inducible and dependent mechanism targets the induction of IFN by inhibition of IRF3 dimerization. This phenomenon is likely to add to the inhibition of SARS-CoV2 induced cytokine release we could observe when using NRF2 agonists. We have previously reported that 4-OI inhibits the expression of STING, which is important for the induction of the IFN-response in cells stimulated with cytosolic DNA(*12*). In line, 4-OI inhibited the IFN-response to HSV1 infection and to stimulation with STING agonists dsDNA and cGAMP **(Fig. 4R-T)**.

## Discussion

Altogether, this study demonstrated that the expression of NRF2 dependent anti-oxidant genes was significantly inhibited in COVID-19 patients, and that the NRF2 agonists 4-OI and DMF inhibited both SARS-CoV2 replication, as well as the expression of associated inflammatory markers. The ability of these NRF2 inducers to also reduce potentially pathogenic IFN-and inflammatory responses while retaining their anti-viral properties is unique to these compounds and promotes their applicability to prevent virus-induced pathology. That NRF2 might be a natural regulator of IFN-responses in the airway epithelium is supported by a recent report demonstrating that NRF2 activity is high while IFN activity is low in the bronchial epithelium(*29*). As DMF is currently used as an anti-inflammatory drug in relapsing-remitting MS, this drug could be easily repurposed and tested in clinical trials to test its ability to limit SARS-CoV2 replication and inflammation-induced pathology in COVID-19 patients. Our observation that 4-OI strongly inhibits the IFN-response to both cytosolic DNA and d RNA, which are canonical anti-viral pathways, but still retain its ability to block viral replication also suggests a spectrum of unidentified cellular programs that are inducible through NRF2 and efficiently restrict viral replication independently of IFNs. This is supported by already mentioned negative correlation between expression of NRF2-inducible genes and infection with HSV1 discovered by Wyler *et al*., through single cell transcriptome analysis(*13*). Similarly to how IFNs block viral replication through the induction of hundreds of effector IFN-stimulated effector genes, the NRF2 controlled anti-viral program might also consists of a myriad of mechanisms that restrict viral replication each by targeting distinct stages of viral replication.

Future studies will determine if patients developing severe SARS-CoV2 pathology also have an underlying NRF2 deficiency, leading to reduced control of viral replication, coupled with excess inflammatory responses. Further, it could be valuable to investigate if patients already in DMF therapy have altered susceptibility to SARS-CoV2 infection and if those infected have milder symptoms and reduced cytokine load. Finally, the fact that 4-OI effectively limited replication of several human pathogenic viruses demonstrated that 4-OI, or related chemically modified compounds such as DMF, could be evaluated as broad-spectrum anti-viral agents for protection against seasonal and pandemic viral infections in general.

## Author contributions

DO and CH conceived the project and prepared figures. DO, ALH, JT, JC, BH, MI, EBS, AK, LR, MI, MS, SF, MT, ML, HH, VG, AH, AK, CG, DVDH, CM, LB, AT, TA, JH, RO, and AA performed experiments, analyzed data, and prepared figures. EF performed bioinformatics and prepared figures. TM was responsible for including COVID-19 patients and achieving patient material. TA, JR, AA, TM, SP, and CH planned experiments and analyzed data. CH and DO drafted and finalized the manuscript. JH planned experiments and edited the manuscript. EF, SP, SB and ML performed bioinformatics and prepared figures. MJ facilitated SARS-CoV2 laboratories. Authors declare no competing interests.

## Data and materials availability

All data is available in the main text or the supplementary materials,

