## Supplementary M & M and figures for "Identification of SARS-CoV2-mediated suppression of NRF2 signaling reveals a potent antiviral and anti-inflammatory activity of 4-octyl-itaconate and dimethyl fumarate"

**This PDF file includes:**

Materials and Methods

Figs. S1 to S10.

### Materials and Methods

Cell lines, reagents and culture conditions

Human lung adenocarcinoma epithelial A549 cells, immortalized human HaCaT keratinocytes, Calu-3 epithelial lung cancer cells and human embryonic kidney HEK293T cells were kindly provided by Søren R. Paludan (Aarhus University, Denmark) and cultured in DMEM (Lonza) supplemented with 10% heat inactivated fetal calf serum, 200 $IU.mL^{-1}$ penicillin, 100 $\mu g.mL^{-1}$ streptomycin and 600 $\mu g.mL^{-1}$ glutamine (hereafter termed DMEM complete). Vero E6 cells expressing hTMPRSS2 were a kind gift of Makoto Takeda (University of Tokyo, Japan)(*1*) and were cultured in DMEM (Lonza) supplemented with 10% heat inactivated fetal calf serum, 200 $IU.mL^{-1}$ penicillin, 100 $\mu g.mL^{-1}$ streptomycin, 600 $\mu g.mL^{-1}$ glutamine and 10 $\mu g.mL^{-1}$ blasticidin. All cell lines were regularly tested for mycoplasma contamination by sequencing from GATC Biotech (Germany). 4-octyl-itaconate (4-OI) was chemically synthetized by Thomas B. Poulsen (Aarhus University, Denmark) and was dissolved in DMSO as previously described here(*2*).

To obtain bone marrow-derived cells (BMDCs), a cell suspension from femurs of C57BL/6 mice (Charles River) was cultured for 6-8 days at 37ºC in RPMI-1640 medium supplemented with 3% FBS and 10% of J558 cell line supernatant containing GM-CSF. Cells were seeded at a density of $10^6$ cells/ml and medium was partially replaced every 2 days. BSC-1 cells (ECACC) were maintained in DMEM supplemented with 5% FBS, 2 mM glutamine, 200 IU/ml penicillin and 100 IU/ml streptomycin at 37ºC 5% CO2.

For generation of KO cell line clones in HaCaT cells specific guide RNA sequences targeting STING (5'-AGAGCACACTCTCCGGTACC-3') or STAT1 (5'-

TTAATGATGAACTAGTGGAG-3') were cloned into the plasmids pX461 (Addgene) (STING) or LentiCRISPR v2 (Addgene) (STAT1). Wildtype HaCaT cells were transfected with the plasmids using the Lipofectamine 2000 Reagent (Invitrogen, Life Tecnologies). 72 h post transfection, the GFP expressing cells were sorted as single cells by FACS and clones were grown to larger cultures (STING). Or 24 h post transfection, the cells were seeded in a dilution sufficient to obtain single cells clones after the puromycin selection. The 2 μg $mL^{-1}$ puromycin selection was initiated 48 h post transfection and continued for 72 h (STAT1). Hereafter, single cell clones were grown to larger cultures which were validated for absence of protein by western blotting and functional analysis to confirm the biological effect of the gene deficiency.

Viruses

We used the SARS-CoV2 strain #291.3 FR-4286 isolated from a patient in Germany, and kindly donated by professor Georg Kochs (Freiburg). The virus was propagated in Vero-TMPRSS2 cells(*3*). Validation and SARS-CoV2 genome detection was performed with Taqman based qPCR using SARS-CoV2 specific primers and probes with the following sequences: Forward primer: AAATTTTGGGGACCAGGAAC, reverse primer: TGGCACCTGTGTAGGTCAAC, Probe: FAM-ATGTCGCGCATTGGCATGGA-BHQ. HSV-1 KOS strain expressing GFP (HSV-1–GFP), HSV-2 333 strain and HSV-2 MS strain were kindly provided by Søren R. Paludan (Aarhus University, Aarhus, Denmark). All HSVs were propagated in Vero cells, purified by ultra-centrifugation, and titrated by standard plaque assay as previously described (*4*). HaCat cells were infected with the different HSVs at a multiplicity of infection (MOI) of 0.01 in a small volume of serum-free medium for 1 h at 37°C. Prior to analysis, cells were incubated with DMEM complete for an additional day of culture. VACV Western reserve strain (VACV-

WR) was a recombinant vaccinia virus (VACV) named vtag2GFP expressing the tag2GFP under the control of strong synthetic VACV early/late promoter was kindly provided by Dr. Rafael Blasco (INIA, Spain). VACV-WR and vtag2GFP stocks were semi-purified by centrifugation through a 36% sucrose cushion and titrated twice by plaque assay. The Brazilian ZIKV isolate ZIKV/H.sapiens/Brazil/PE243/2015 was originally described in (*5*) and was grown on Vero cells. Viral titers were determined by plaque assay on A549 BVDV NPro cells (kind gift from R. Randall, St Andrews). These cells are optimized for virus growth as they stably express the NPro protein of bovine viral diarrhea virus (BVDV), which induces degradation of IRF3 (*6*).

<u>Viral entry assay</u>

Quantification of HSV-1 entry in the presence of 4-OI was performed using the cold binding assay previously described(*7*). Cells were pretreated with 4-OI (150 μM) or DMSO (control) for 48 hours. Cells were pre-incubated at 4°C for 30 minutes, then incubated with HSV-1 at a MOI of 10 for 1 hours at 4°C. Cells were then shifted to 37°C for 1 hours to activate virus internalization. After that, cells were washed twice with PBS, then uninternalized virus particles were washed with citric acid buffer (135 mM NaCl, 10 mM KCl, 40 mM citric acid, pH 3) incubation for 5 minutes, then cells were washed twice more with PBS. Cells were scraped and genomic DNA was extracted using QIAamp DNA mini kit (QIAGEN). Quantitative PCR were performed using UL30-F and UL30-R primers for HSV-1 genomic DNA. Primer sequence: UL30-F: ACATCATCAACTTCGACTGG, UL30-R: CTCAGGTCCTTCTTCTTGTCC

Primary cells and culture conditions

Peripheral Blood Mononuclear cells (PBMCs) were isolated from healthy donors (blood donors gave written consent as accordingly to the ethical guidelines at Aarhus University Hospital) by Ficoll Paque gradient centrifugation (GE Healthcare). Monocytes were separated using a monocyte enrichment kit (STEMCELL) according to the manufacturer's instructions or from PBMCs by adherence to plastic in RPMI 1640 supplemented with 10% AB-positive human serum. Differentiation of monocytes to macrophages was achieved by culturing in Dulbecco's Modified Eagle Medium (DMEM) supplemented with 10% heat inactivated AB-positive human serum 9 days in the presence of 10 ng/ml M-CSF (R&D Systems), as previously reported in (*8*).

Patients included in this study

All patients were positive for SARS-CoV2 by PCR from throat swab and admitted to the ICU and receiving ventilatory support due to severe pneumonia with a component of acute respiratory distress syndrome (ARDS). P1 27-years old male, day 14 at in ICU. P2 57-year-old female, day 9 in ICU. P3 43 years-old male, day 5 in ICU.  P4 28 years-old female, day 3 in ICU.

Air-Liquid Interface Epithelium model

Primary nasal cells were isolated using a nasal brush (Dent-O-Care, #620B) inserted into the nasal turbinations and twisted. Cells were isolated from the brush by gently expelling monolayer medium (Airway Epithelial Cells Basal Medium, PromoCell, #C-21260 + 1 pack of Airway Epithelial Cell Growth Medium Supplement, PromoCell, #39160 + 100 U/ml Penicillin/Streptomycin, Gibco #10378) and PBS to wash of cells. Cells were cultured in monolayer culture in tissue culture flask (Sarstedt, TC: Standart #83.3911) coated with 0,1 mg/ml Bovine type I collagen solution (Sigma-Aldrich, #804592, diluted in sterile $ddH_2O$). Monolayer cultures were split using 1x Trypsin mixed with 0.3 mM EDTA (10x Trypsin (2,5 %), Gibco, #15090, diluted to working concentration in PBS + UltraPure 0.5 M EDTA, Invitrogen, #15575) at approx. 80% confluency. At passage two, cells were seeded at 2-3x10^4 cells on 6,5 mm Transwell membranes (Corning, #3470) coated with 30 ug/ml Bovine type I collagen solution (Sigma-Aldrich, #804592, diluted in sterile $ddH_2O$). Cells were seeded and submerged in 2x P/S (200 U/ml Pen/Strep) DMEM-low glycose (Sigma-Aldrich, D5921) mixed one to one with 2x Monolayer medium (Airway Epithelium Cell Basal Medium, (PromoCell, #C-21260) supplemented with 2 packs of Airway Epithelial Cell Growth Medium Supplement (PromoCell, #C-39160) without triiodothyronine + 1 ml of 1.5 mg/ml BSA). When cultures reach full confluency ALI (=Air-liquid interface) is introduced and medium is changed to ALI medium (Pneumacult ALI medium kit (StemCell, #5001) with ALI medium supplement (StemCell, #5001) and 100 U/ml Pen/strep) supplemented with 24 ug of hydrocortisone (StemCell, #07925) and 0.2 mg heparin (StemCell, #07980). Membranes was allowed at least 21 days of differentiation verified by extensive cilia beating and mucus covering.

Upon initiation of treatment ALI cultures was washed for 5 minutes using DMEM (low glycose, no additives) and baso-lateral medium changed for ALI medium containing either 150 uM 4-OI or DMSO. Baso-lateral medium containing treatment was left overnight. 100 ul of DMEM (low glycose) with 150 uM or DMSO was additionally added to the apical compartment overnight. At time of infection, apical medium was removed and 100 ul DMEM (low glycose) containing SARS-CoV-2 at MOI 0.1 was added to all membranes for 1 hour and placed in 37 $^{o}$C incubator. After 1-hour apical infection medium was removed and membranes placed in 37 $^{o}$C incubator for 24 hours before harvest.

At time of harvest, baso-lateral medium was removed and 500 ul Trypsin/EDTA was added bao-laterally and 200 ul was added apically. After approx. 5 minutes cells were harvested using 5% FPB/DMEM (low glycose). Cells were lysed for RNA isolation using lysisbuffer from High Pure RNA Isolation Kit (Roche Diagnostics, #11828665001). For Western blot cells were lysed in RIPA buffer containing 1/10 Protease inhibitor (Roche), 1/1000 Benzonase (Sigma, #E1014) and 1/50 0.5 M Sodium Flouride.

Short-interfering RNA (siRNA)-mediated knock down

For short interfering RNA experiments, HaCat cells were transfected in 6-well plates with 80 pmol of human Nrf2(1) (sc-37030) or control si RNA (sc-37007) diluted in serum and antibiotic free DMEM and using Lipofectamine RNAi Max as per manufacturer's instructions. HaCat cells were incubated for 72h in the presence of the siRNA before being processed. For experiments in Calu-3 cells, the same protocol conditions were used with control si RNA (sc-37007), Keap1 siRNA (sc-43878) and HO-1 (sc-35554) that were lipofected for 48h.

dsDNA, cGAMP and optimized RIG-I agonist stimulation of cells.

HSV-60 naked, a viral dsDNA motif, 2'3'-cGAMP, a STING ligand, and M8, a sequence optimized RIG-I agonist (*9*) were obtained from Invivogen and John Hiscott (Pasteur Institute, Rome), respectively. Intracellular delivery of dsDNA and cGAMP was achieved using Lipofectamine 2000 (Invitrogen) diluted in serum-free medium with a ratio of Lipo.dsDNA/cGAMP of 1:1. Final concentration for both dsDNA and cGAMP was $4\mu g.mL^{-1}$. Intracellular delivery of M8 was achieved using Lipofectamine RNAiMax (Invitrogen) diluted in serum-free medium with a ratio of Lipo.RNA of 1:1. Final concentration of M8 was $10\ ng.mL^{-1}$.

VACV infection assays.

BMDCs, HaCaT and HaCaT Stat1 KO were incubated or not with 150 µM 4-OI for 48 h before infection with vtag2GFP using 0.1 or 0.01 pfu/cell at 37 ºC for 60 min. Then, infected cells were washed to remove potential unbound viruses and infection proceeded at 37 ºC. To determine the proportion of vtag2GFP infected cells GFP expression was detected at 16 hpi by flow cytometry using triplicates. Briefly, cells were harvested, washed with FACS buffer (PBS, 0.01% sodium azide, and 0.1% BSA) and fixed with paraformaldehyde 4% in PBS for 10 min. After extensive washing with FACS buffer, $2x10^4$ cells were scored and analyzed in a FACSCalibur flow cytometer (BD Sciences) per experimental condition in triplicates. To determine VACV virus titres, HaCaT and HaCaT STAT1 KO cells were previously stimulated for 48 h or not with 150 µM 4-OI and then infected with VACV-WR using 0.1 or 0.01 pfu/cell at 37 ºC. At 24 hpi, cells were harvested in their own media, centrifuged at 1,800 × g for 5 min, and resuspended in 0.5 ml of fresh medium. In all cases, samples were frozen, thawed three times and titrated using duplicates in BSC-1 cells. Briefly, preconfluent monolayers of BSC-1 cells were infected with 10-fold serial dilutions of viral inoculums for 1h at 37ºC. Then, inoculum was replaced with semi-solid carboxy-methyl cellulose (Sigma) media with 2 % FBS and cells fixed in 10 % formaldehyde at 3 dpi. Plaques were stained with 0.1% (w/v) crystal-violet. Two independent experiments were performed.

Zika Virus infections.

A549 cells (kind gift from G. Kochs, Freiburg) and Huh-7 were cultured at 37°C in Dulbecco's Modified Eagle Medium (DMEM), supplemented with 10% FCS and 2mM L-Glutamine. Cells were seeded in 24 well plates and pre-treated with 4-octyl-itaconate (4-OI) (150uM) for 48h. Cells were infected with ZIKV (moi 0.1) for 1h. 4-OI was freshly added when the medium was changed.Cells were lysed and total RNA was extracted at 96hpi using the QIAshredder (Qiagen) and RNeasy Mini Kit (Qiagen) according to the manufacturer's instructions. RNA was reverse transcribed using SuperScript II Reverse Transcriptase (Invitrogen) into cDNA that was then used for qPCR with SYBR green PCR kit (Life Technologies). $C_T$ values were normalized to GAPDH ($\Delta C_T$). SYBR green primer probes used include GAPDH (for: CATGGCCTTCCGTGTTCCTA, rev: CCTGCTTCACCACCTTCTTGA) and ZIKV (for: CGAGGAACATCCAGACTC, rev: ATTGGAGATCCTGAAGTTCC).

SARS-CoV2 TCDI50% assay

The assay was performed as follows. 2 x $10^4$ Vero E6 TMPRSS2 cells were seeded in 90ul DMEM (Gibco, + 2% FCS (Sigma-aldrich) + 1% Pen/Strep (Gibco) + L-Glutamine (Sigma-Aldrich) per well in flat-bottom 96-well plates. 24h after, samples were titrated onto the cells by addition of 10ul of a 10-fold serial dilution. One full plate was used per sample analyzed. Each dilution of supernatant were represented 8 times on a plate. The cells were incubated for 72h in a humidified $CO_2$ incubator at 37 ˚C, 5% $CO_2$, before fixing with 5% Formalin (Sigma-Aldrich) and staining with crystal violet solution (Sigma-Aldrich). Images were taken using a Leica DMi1, microscope with a Leica MC170 HD camera. TCDI50 % virus titer calculated by Reed-Muench method.

Western blot.

Cells were lysed in 100 µL of ice-cold Pierce RIPA lysis buffer (Thermo Scientific) supplemented with 10 mM NaF, 1x complete protease cocktail inhibitor (Roche) and 5 $IU.mL^{-1}$ benzonaze (Sigma), respectively. Protein concentration was determined using a BCA protein assay kit (Thermo Scientific). Whole-cell lysates were denatured for 3 min at 95°C in presence of 1x XT Sample Buffer (BioRad) and 1x XT reducing agent (BioRad). 10-40 µg of reduced samples was separated by SDS-PAGE on 4-20% Criterion TGX precast gradient gels (BioRad). Each gel was run initially for 15 min at 70V and 45 min at 120V. Transfer onto PVDF membranes (BioRad) was done using a Trans-Blot Turbo Transfer system for 7 min. Membranes were blocked for 1h with 5% skim-milk (Sigma Aldrich) at room temperature in PBS supplemented with 0.05% Tween-20 (PBST). Membranes were fractionated in smaller pieces and probed overnight at 4°C with any of the following specific primary antibodies in PBST: anti-Nrf2 (12721, Cell Signaling 1:1000), anti-TBK1/NAK (3013, Cell Signaling 1:1000), anti-phospho-TBK1/NAK (5483, Cell Signaling 1:1000), anti-SQSTM1/p62 (8025, Cell Signaling 1:1000), anti-IRF3 (11904, Cell Signaling 1:1000), anti-phospho-IRF3 (4947, Cell Signaling 1:500), anti-HO-1 (5853, Cell Signaling 1:1000), anti-IFIT1 (14769, Cell Signaling 1:1000), anti-NRF2 (12721, Cell Signaling 1:1000), anti-STING (13647, Cell Signaling 1:1000), anti-NqO1 (3187, Cell Signaling 1:1000), and anti-Vinculin (18799, Cell Signaling 1:1000) used as loading control. After three washes in PBST, secondary antibodies, peroxidase-conjugated F(ab)2 donkey anti-mouse IgG (H+L) (1:10000) or peroxidase-conjugated F(ab)2 donkey anti-rabbit IgG (H+L) (1:10000) (Jackson ImmunoResearch) were added to the membrane in PBST 1% milk for 1h at room temperature. All membranes were washed three times and exposed using either the SuperSignal West Pico PLUS chemiluminescent substrate or the SuperSignal West

Femto maximum sensitivity substrate (ThermoScientific) and an Image Quant LAS4000 mini imager (GE Healthcare).

Semi-native WB Dimerization assay.

IRF3 dimerization was assayed under semi-native conditions. Cells were lysed in ice-cold Pierce RIPA lysis buffer (Thermo Scientific) supplemented with 10 mM NaF, 1x complete protease cocktail inhibitor (Roche) and 5 IU/mL benzonaze (Sigma). Protein concentration was determined using a BCA protein assay kit (Thermo Scientific). Whole-cell lysates were mixed with 1x XT Sample Buffer (BioRad); samples were neither reduced nor heated before separation was done on 4-20% Criterion TGX precast gradient gels (BioRad) by SDS-PAGE electrophoresis. Each gel was run initially for 15 min at 70V and 15 min at 120V. Transfer onto PVDF membranes (BioRad) was done using a Trans-Blot Turbo Transfer system for 7 min. Membranes were blocked for 1h with 5% skim-milk (Sigma Aldrich) at room temperature in PBS supplemented with 0.05% Tween-20 (PBST). Membranes were probed overnight at 4°C with the following specific primary antibody in PBST: anti-IRF3. After three washes in PBST, secondary antibodies, peroxidase-conjugated F(ab)2 donkey anti-rabbit IgG (H+L) (1:10000) (Jackson Immuno Research) were added to the membrane in PBST 1% milk for 1h at room temperature. All membranes were washed three times and exposed using either the SuperSignal West Pico PLUS chemiluminescent substrate or the SuperSignal West Femto maximum sensitivity substrate (Thermo Scientific).

qPCR analysis.

Gene expression was determined by real-time quantitative PCR, using TaqMan detection systems (Applied Biosciences). RNA was extracted using the High Pure RNA Isolation kit (Roche) and RNA quality was assessed by Nanodrop spectrometry (Thermo Fisher). RNA levels were analyzed using premade TaqMan assays and the RNA-to-Ct-1-Step kit according to the manufacturer's recommendations (Applied Biosciences).

Transcriptome analysis COVID19

COVID19 data set analysis (Fig. 1): RNA-seq data was obtained from an already available dataset from Blanco-Melo (doi: https://doi.org/10.1101/2020.03.24.004655). From the raw read-counts differential expression values were calculated using DESeq2(*10*). Significantly differentially expressed genes (SDEGs) were selected based on the thresholds of adjusted P-value <0.05 and absolute fold change of 2 (Fig. 1A). In order to focus on commonality across the different conditions with respect to generating clinically relevant hypotheses, the 815 SDEGs obtained from the biopsy sample were checked if these were also present in the list of SDEGs in the other conditions. All genes that occur in at least 3 or more other conditions were included in the final list, resulting in 113 genes. Finally, the expression values for all genes in the final list across all conditions were assembled, clustered using Euclidean distance metric and Ward's variance minimization algorithm, and visualized as a heatmap using Python3.7 and seaborn cluster-map tools. Then the gene-sets from each of the outlined clusters were used for pathway enrichment analysis using Enrichr (Fig. 1B) (*11*). Finally, the STRING database(*12*) was used to construct the cloud network starting from lists of genes manually annotated for NRF2, inflammation and IFN signaling. Edges and nodes were extracted from STRING and imported to Cytoscape (*13*) version 3.7 for further visualization (Fig. 1C).

Transcriptome analysis of HSV1-infected HaCaT cells

RNA sequencing was performed in collaboration with BGI Europe Genome Center (Copenhagen, Denmark) following the standard operational procedure as described before(*14*). Briefly, the quality of total RNA was checked using the Agilent 2100 bioanalyzer. To construct the sequencing library for MGIseq-2000, approximately 1 μg of polyA enriched RNA was used for library construction using the MGIEasy RNA Directional Library Prep Kit (MGI Tech). Next, paired-end sequencing with 100 cycles was performed using the MGISEQ-2000 sequencing instrument, according to the manufacturer's instructions. We generated an average of 63 million raw reads for each sample. The clean RNA reads were first aligned to the hg19 UCSC RefSeq (RNA sequences, GRCh37) using bowtie2 at first. To map the transcripts from the viruses, the unmapped reads were then aligned to the coding sequence of the human herpesvirus 1 (KOS strain). The expression of human genes and virus genes were performed by transforming mapped transcript reads to TPM using RSEM(*15*). The normalized expression were estimated and normalized by DEseq2. Differentially expressed genes were defined as genes with fold change over two-folds and adjusted P-value less than 0.001 using DESeq2.

Cell Painting Assay

The procedure from Bray(*16*) et al was followed with adaption to 96-well plates and optimized fluorophore concentrations for Vero E6 cells and the Celldiscoverer 7 imaging system:

5000 wt Vero E6 cells were seeded into the inner 60 wells of a 96-well plate with optical bottom (Corning #3603) in 75 µL full growth medium. After 24 h, compounds were dosed as a 25 µL 4X solution (final DMSO = 0.5%). After 24 h, 75 µL medium was removed and replaced with 75 µL medium containing 500 nM MitoTracker Deep Red (final C = 325 nM) and plates were incubated (37 °C, 5% CO2, humid) in the dark for 30 min. Wells were then aspirated and 75 µL medium were added, before adding 25 µL 16% paraformaldehyde (Electron Microscopy Sciences, cat. no.15710- S) (final PFA = 4 %) and incubating in the dark for 20 min. Plates were washed once with 1X HBSS (Invitrogen, #14065-056) and 75 µL 0.1% (vol/vol) Triton X-100 (BDH, #306324N) in 1X HBSS was added and incubated for 15 min in the dark. Plates were washed twice with 1X HBSS before addition of 75 µL multiplex staining solution (final concentrations: 0.75 µg/mL WGA-AF555; 8.75 µg/mL Concanavalin-AF488; 2.5 µL/mL Phalloidin-AF568; 0.75 µM SYTO 14; 5 µg/mL Hoechst 33342 in 1 % (wt/vol) bovine serum albumin (Sigma-Aldrich, #A9647)) and incubation for 30 min in the dark. Plates were then washed three times with 1X HBSS, with no final aspiration and imaged immediately in a Zeiss Celldiscoverer 7 automated microscope. 9 images are acquired in each well with 2x2 binning using the AxioCam 702 CMOS 12-bit camera with 4x analog gain. To generate the bioactivity profiles the workflow outlined in Svenningsen & Poulsen(*17*) was followed. In short, CellProfiler 2.1.1(*18*) was used to correct images for uneven illumination followed by image segmentation and extraction of 1476 features across nuclei, cytoplasm and the whole cell on a per-cell basis. Features were then averaged to per-well profiles after which the data was normalized on a per-plate basis followed by per-treatment aggregation which affords the final profiles. The heatmap of morphological profiles is visualized with heatmap.2 in the gplots package. The Pearson correlation matrix is calculated using the stats package in R 3.6.0 (R core Team) and visualized using the corrplot 0.84 package (R package corrplot).

Hierarchical clustering of the correlation matrix is performed using the stats package with Pearson correlation coefficients as distance metric and average linkage method. To determine activity scores and thresholds the workflow described by Hutz et al.(*19*) was followed.

Data availability

RNA sequencing files generated for analysis of HaCaT cells infected with HSV in the presence or absence of 4-OI have been deposited to the data depository database (CNGBdb, https://db.cngb.org) with the following accession number : CNP0001039.

Ethics

The project was approved by Institutional review boards at Aarhus University Hospital, by the Danish National Committee in Health Research Ethics (1-10-72-80-20) and the Danish Data protection Agency in accordance with the ethical standards of the Helsinki Declaration. Written informed consent was obtained from all study participants.

Statistical analysis.

Values were expressed as the means ± SEM. Graphs and statistics were computed using Graph Pad Prism 7. An unpaired, two-tailed Student's *t*-test was used to determine significance of the difference between the control and each experimental condition. *P* values of less than 0.05 were considered statistically significant, ***, $p<0.001$; **, $p<0.01$, and *, $p<0.05$.

**Fig. S1.**

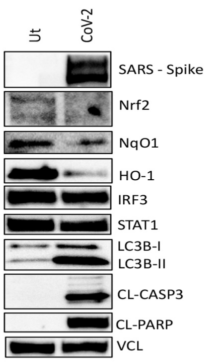


**Fig. S1.** NRF2 inducible genes are repressed during infection with SARS-CoV2 in Vero cells. Vero cells were either left untreated or infected with SARS-CoV2 for 24 hours before analysis of cell pellets by immunoblotting using indicated antibodies. Experiment is one representative of two independent experiments.

**Fig. S2.**

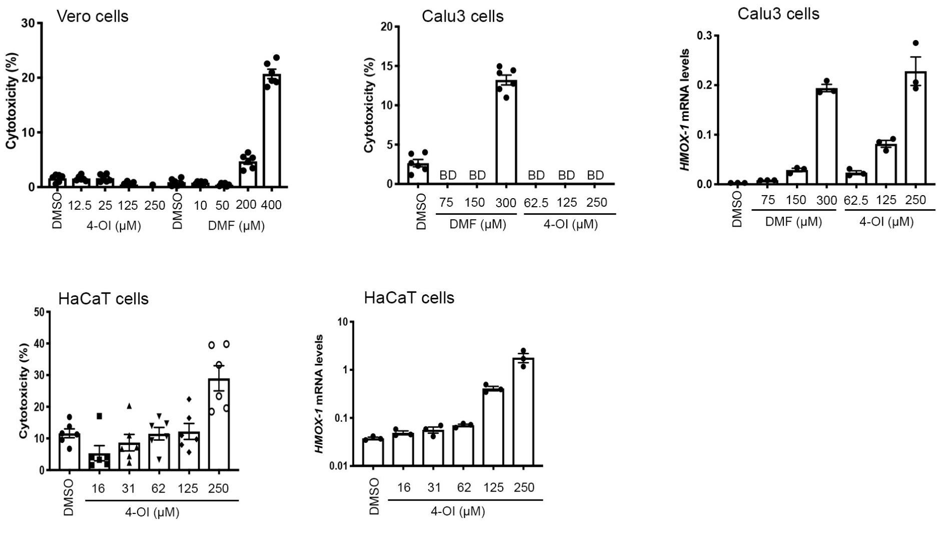


**Fig. S2.** Cytotoxicity of 4-OI and DMF
Indicated cell lines were treated with either 4-OI or DMF for 48 hours before analysis of cell supernatants by LDH assay and of cell pellets by qPCR for NRF2-inducible gene HO-1 (*HMOX-1*).

**Fig. S3.**

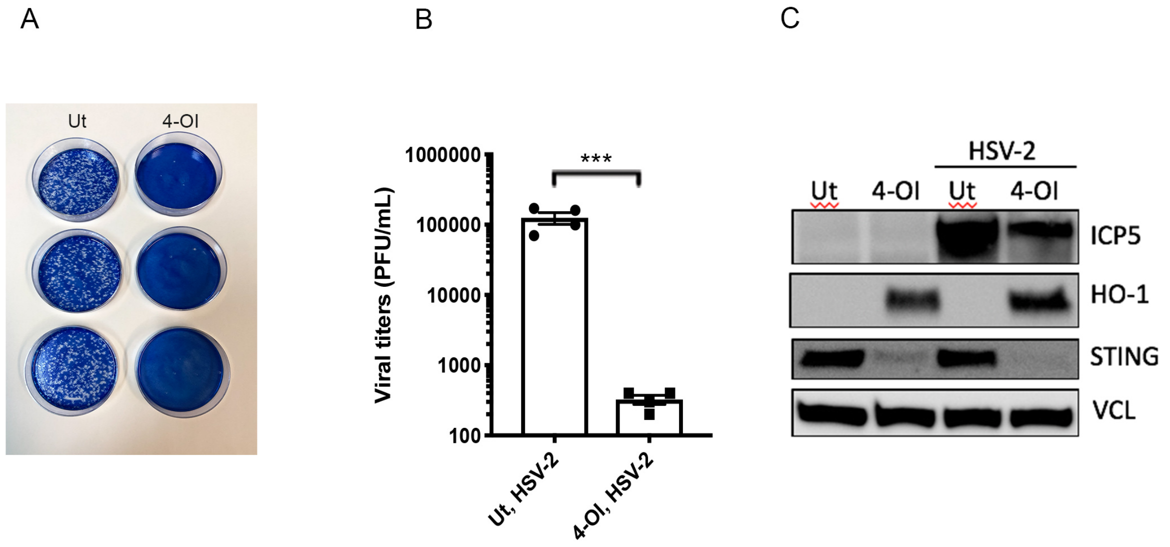


**Fig. S3.** 4-OI inhibits HSV2 replication
HaCaT cells were treated with 4-OI at 125μM for 48 hours before infection with HSV2 at MOI 0.01. Cell supernatants were then harvested to determine release of progeny virus by plaque assay (A+B) and cell pellets were lysed for immunoblotting using specific antibodies against HSV protein ICP5, Heme Oxygenase 1 (HO-1), STING, and Vinculin as loading control(C). A) is a photo of plaque assay performed on vero cells. B) is a quantification of the plaque assay. Experiements are representative of two independent experiments. Statistical analysis was performed using a two-tailed Student's t-test where ***$p<0.001$.

**Fig. S4.**

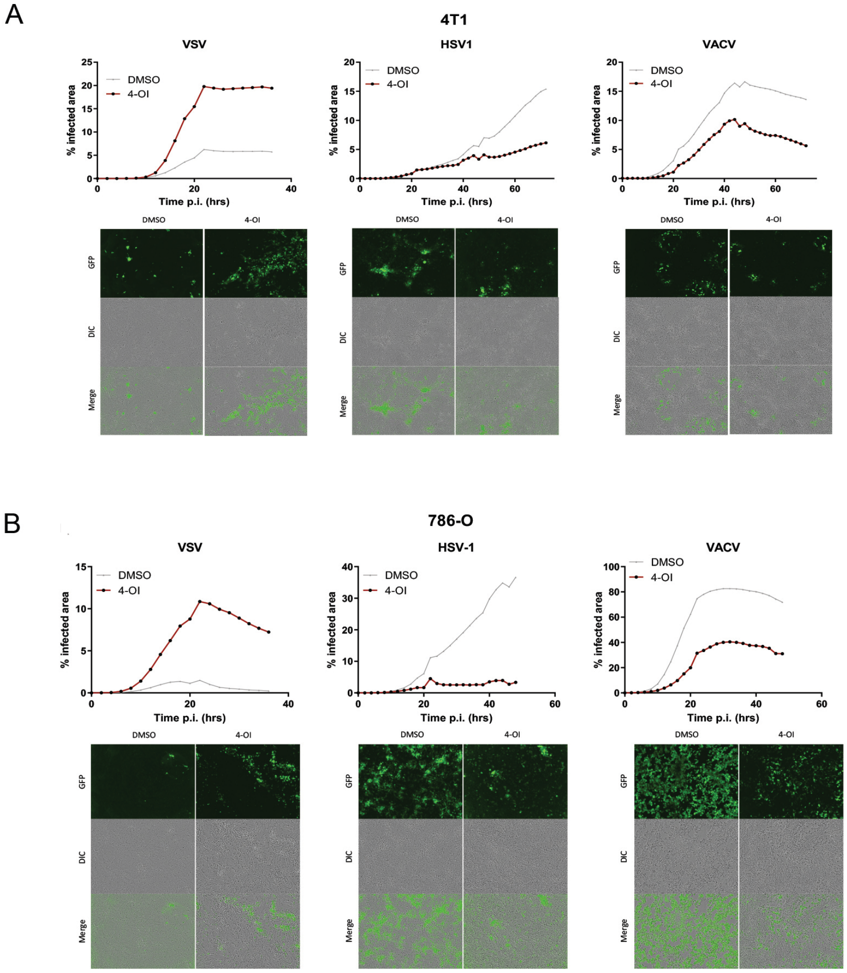


**Fig. S4**. 4-OI impairs HSV and VACV spread in cancer cells but enhances VSV.
(A) Mouse breast cancer cells (4T1) or (B) human renal carcinoma cells (786-O) were treated with 4-OI (150 μM) for 48 hours, then infected with VSV-Δ51-GFP, HSV-ΔICP0-GFP or VACV-GFP at MOI of 0.1. Upper line charts: Live virus spreads were monitored via GFP fluorescence signal using the Incucyte Live-Cell imaging system, images were taken very 2 hours. Lower images: representative fluorescence microscopy images of virus infection at 24 hours p.i. for VSV and 48 hours p.i. for HSV-1 and VACV. Data are representative of two independent experiments.

### Fig. S5.

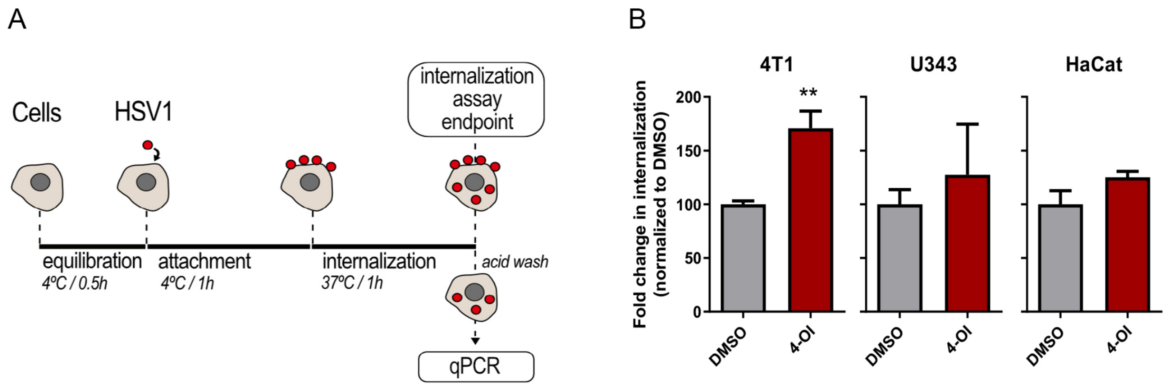


**Fig. S5**. 4-OI enhances HSV-1 internalization. (A) Schematic representation of the internalization assay performed in a breast cancer cell line (4TI), in human glioblastoma (U343), and in immortalized human keratinocyte (HaCaT). Cells were pre-treated with 4-OI (150μM) for 48 hours, then the viral entry assay was carried out with HSV-1 incubation at an MOI of 10. Bars indicate mean with SEM from at least 3 independent experiments. Statistical analysis was performed using unpaired student t-test, **:p<0.01.

### Fig. S6.

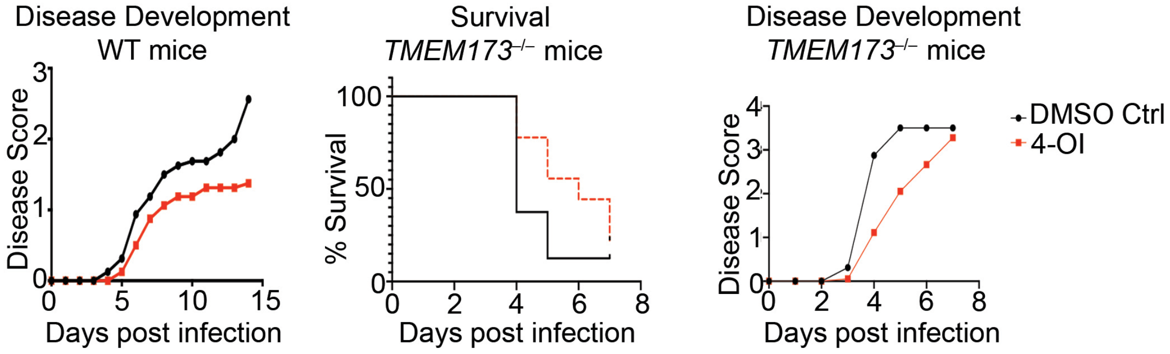


**Fig. S6.** *In vivo* anti-viral effect of 4-OI in model of HSV2
Eight week old wt or *TMEM173*$^{-/-}$ C57BL6J mice were treated intravaginally with 4-OI at 150μM before intravaginal inoculation with HSV2 (333 strain) 24 hours later. Disease progression and survival was monitored and scored for 15 days for WT and 8 days for *TMEM173*$^{-/-}$ mice. Data are representative of two independent experiments with 8 mice (n=8) in each of the experimental groups.

**Fig. S7.**

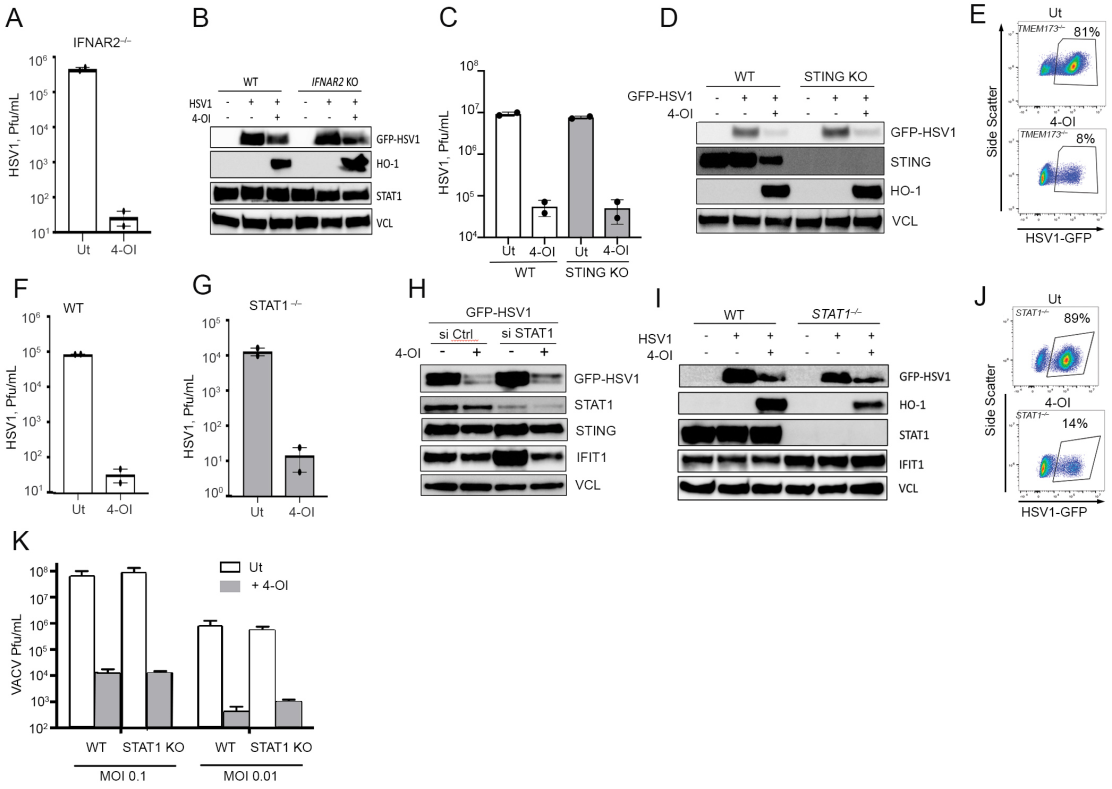


**Fig. S7.** 4-OI induces an anti-viral program independently of IFNs but dependent on NRF2.
**A-K)** WT HaCaT cells or HaCaT cells deficient in IFNAR2, STING or STAT1were either treated with DMSO alone (Ut) or with 4-OI at 125μM before being infected with GFP-expressing HSV1 (GFP-HSV1) at MOI 0.01. After 20 hours, supernatants were collected for analysis by plaque assay (A, C, F, and G). In parallell experiment, cells were either lysed for immunoblotting (B, D, H and I) using indicated antibodies or analyzed by flow cytometry (E and J).
**K)** Cells were treated with 4-OI at 150μM before infection with VACV WR strain at MOI 0.01 and 0.1. After 20 hours, supernatants were collected and analyzed by plaque assay. Data are representative of a minimum of two independent experiments with bars indicating mean +/-s.e.m. of at least two biological replicates.

**Fig. S8.**

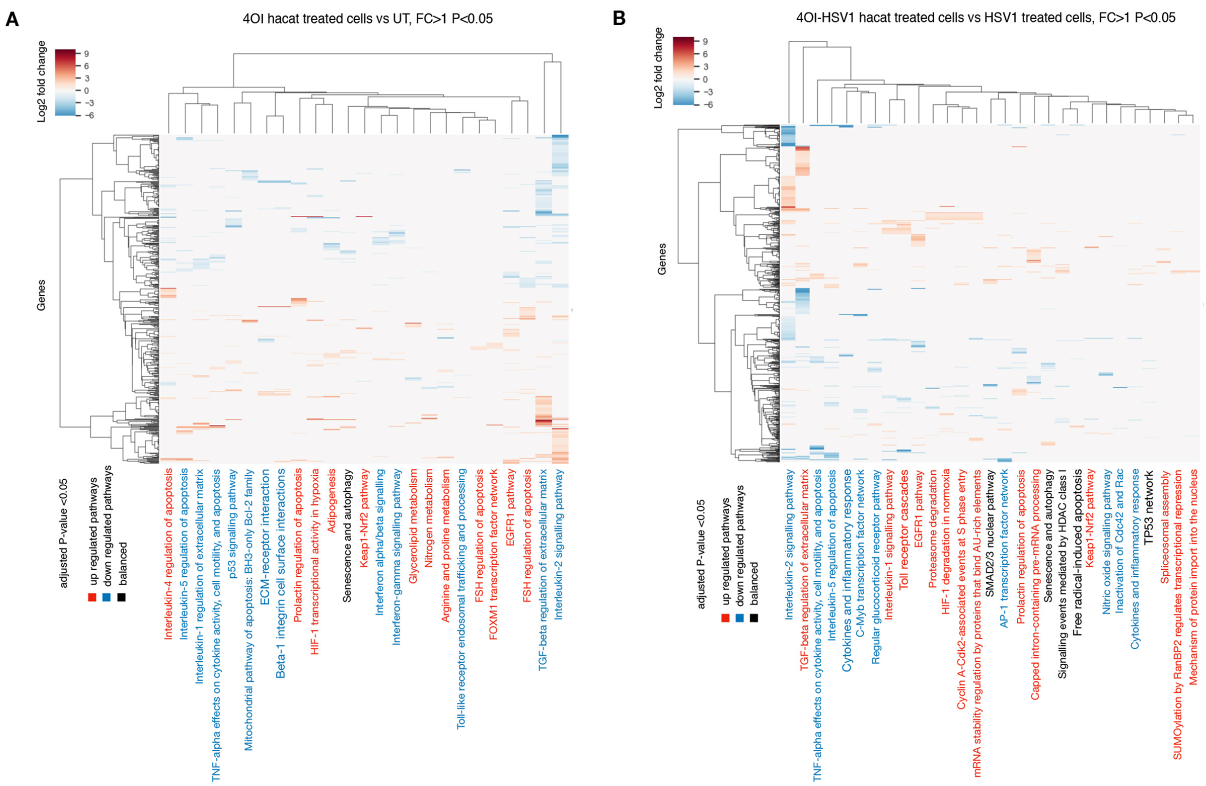


**Fig S8.** Transcriptome pathway analysis of 4-OI treated HaCaT cells
Disregulated pathways significant at adjusted P-value<0.05 in HACAT cells after 4OI treatment relative to un-treated (UT) cells (A) and in HACAT cells after 4OI and HSV1 exposure relative to HSV1 infected cells (B). Both heat maps indicate the presence of a gene involved in each pathway by the genes colour coded expression value. Genes are filtered for |log 2(FoldChange)|>1 and adjusted p-value<0.05. Names of pathways with a higher number of up-regulated genes are shown in red, those with a higher number of down-regulated genes in blue,and are written in black if the number of up- and down-regulated genes are equal.

**Fig. S9.**

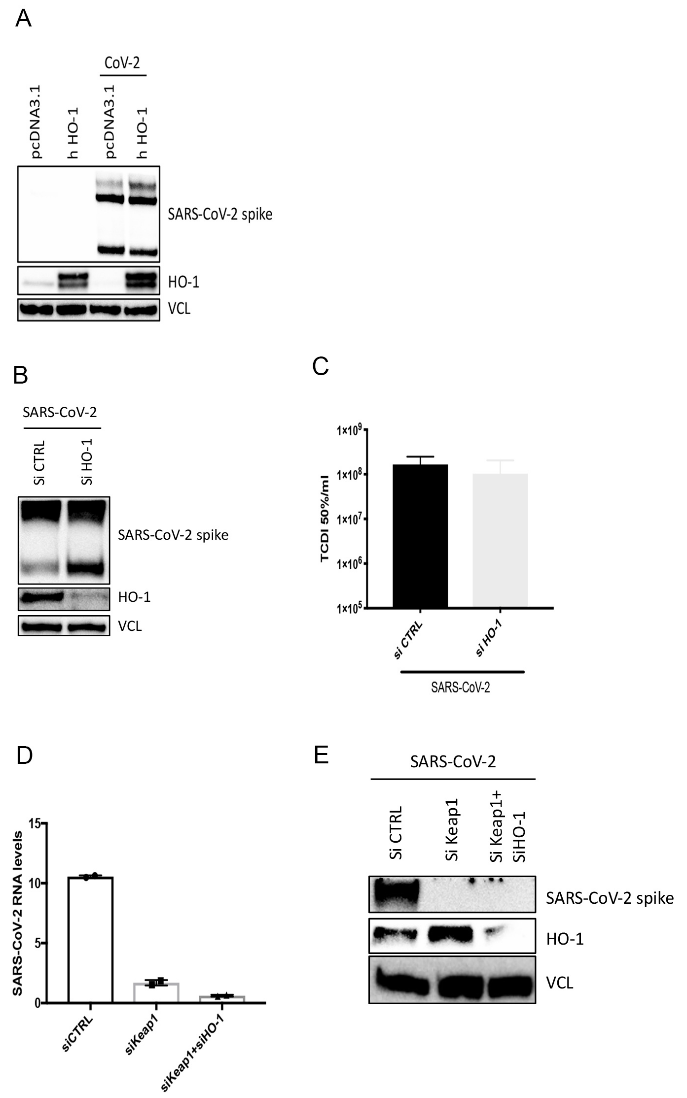


**Fig. S9.** HO-1 is not an important restriction factor for SARS-CoV2
A) Vero cells were transfected with either an empty expression vector or one that encoded expression of human HO-1. Cells were then infected with SARS-CoV2 for 48 hours before analysis by immunoblotting. Data are one representative of two independent experiments.
B+C) Calu-3 cells were treated with ctrl siRNA or siRNA targeting HO-1. Cells were then infected with SARS-CoV2 for 48 hours before analysis of cell pellet by immunoblotting and of cell supernatants by TCID50 analysis.Data are one representative of two independent experiments.
D+E) Calu-3 cells were treated with indicated siRNAs before infection with SARS-CoV2 for 48 hours. Cell pellets were then analysed by qPCR or by immunoblotting.Data are one representative of two independent experiments.

### Fig. S10

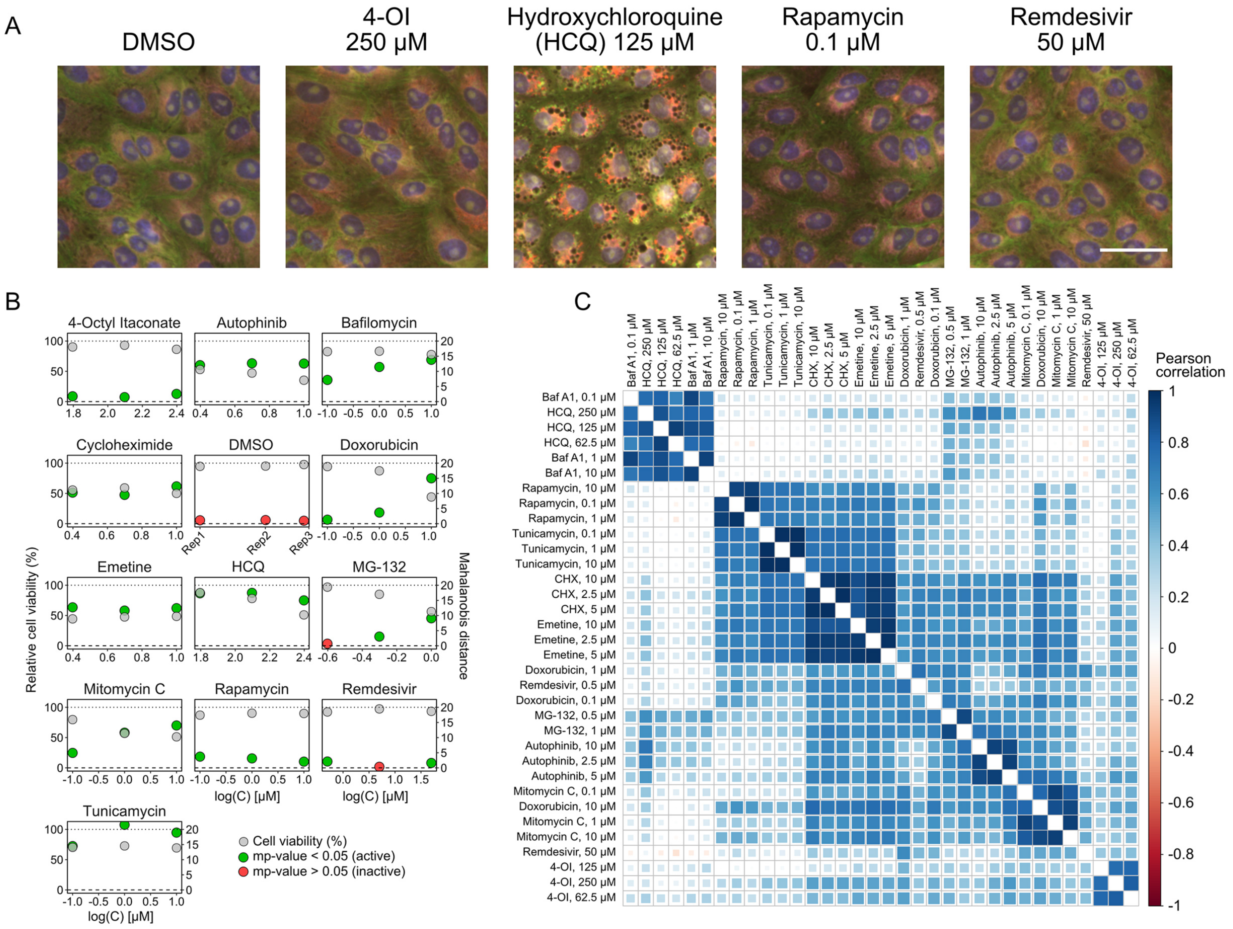


**Fig. S10** Morphological profiling in Vero cells.
Vero E6 cells were subjected to morphological profiling. Compounds were dosed at three concentrations, either 2-fold or 10-fold dilutions in quadruplicates for 24 hours before staining with MitoTracker Deep Red for 30 minutes. The cells were then fixed and permeabilized before staining with the multiplex staining solution and imaged in 5 channels. Image analysis in Cell Profiler 2.1.1 extracted 1476 features related to cell morphology, which constitutes the morphological profile.
A) Representative images from the profiling experiment. Scale bar = 50 µM.
B) Cell viability and profile activity measured as the Mahalanobis distance to DMSO negative controls. The mp-value (see method section) is used to determine significant activity. 4-OI has low but significant activity (mp-value < 0.05) without loss of cell viability. DMSO is quadruplicates of wells kept separate from the DMSO wells used for normalization.
C) Correlation matrix of Pearson correlations clustered by hierarchical clustering with average linkage for active profiles (mp-value < 0.05). The three doses of 4-OI clusters together but show no large correlations to other modulators of various cell processes such as translation (cycloheximide (CHX), emetine), transcription (doxorubicin, mitomycin c) or lysosomal acidification (bafilomycin A1 (Baf A1), hydroxychloroquine (HCQ)).
